# COVID-19 Causes Ciliary Dysfunction as Demonstrated by Human Intranasal Micro-Optical Coherence Tomography Imaging

**DOI:** 10.1101/2022.07.08.499336

**Authors:** Kadambari Vijaykumar, Hui Min Leung, Amilcar Barrios, Courtney M. Fernandez-Petty, George M. Solomon, Heather Y. Hathorne, Justin D. Wade, Kathryn Monroe, Katie Brand Slaten, Qian Li, Sixto M. Leal, Derek B. Moates, Hannah M. Pierce, Kristian R. Olson, Paul Currier, Sam Foster, Doug Marsden, Guillermo J. Tearney, Steven M. Rowe

**Author notes:** **Corresponding author:** Steven M. Rowe, MD MSPH, Address: 1918 University Boulevard, MCLM 824, Birmingham AL 35294. indicates equal contribution (Kadambari Vijaykumar, Hui Min Leung). indicates shared senior authorship (Guillermo J. Tearney, Steven M. Rowe).

## Abstract

Severe acute respiratory syndrome coronavirus (SARS-CoV-2), causative agent of coronavirus disease 2019 (COVID-19), binds via ACE2 receptors, highly expressed in ciliated cells of the nasal epithelium. Micro-optical coherence tomography (μOCT) is a minimally invasive intranasal imaging technique that can determine cellular and functional dynamics of respiratory epithelia at 1-μm resolution, enabling real time visualization and quantification of epithelial anatomy, ciliary motion, and mucus transport. We hypothesized that respiratory epithelial cell dysfunction in COVID-19 will manifest as reduced ciliated cell function and mucociliary abnormalities, features readily visualized by μOCT. Symptomatic outpatients with SARS-CoV-2 aged ≥ 18 years were recruited within 14 days of symptom onset. Data was interpreted for subjects with COVID-19 (n=13) in comparison to healthy controls (n=8). Significant reduction in functional cilia, diminished ciliary beat frequency, and abnormal ciliary activity were evident. Other abnormalities included denuded epithelium, presence of mucus rafts, and increased inflammatory cells. Our results indicate that subjects with mild but symptomatic COVID-19 exhibit functional abnormalities of the respiratory mucosa underscoring the importance of mucociliary health in viral illness and disease transmission. Ciliary imaging enables investigation of early pathogenic mechanisms of COVID-19 and may be useful for evaluating disease progression and therapeutic response.

**Graphical abstract:** 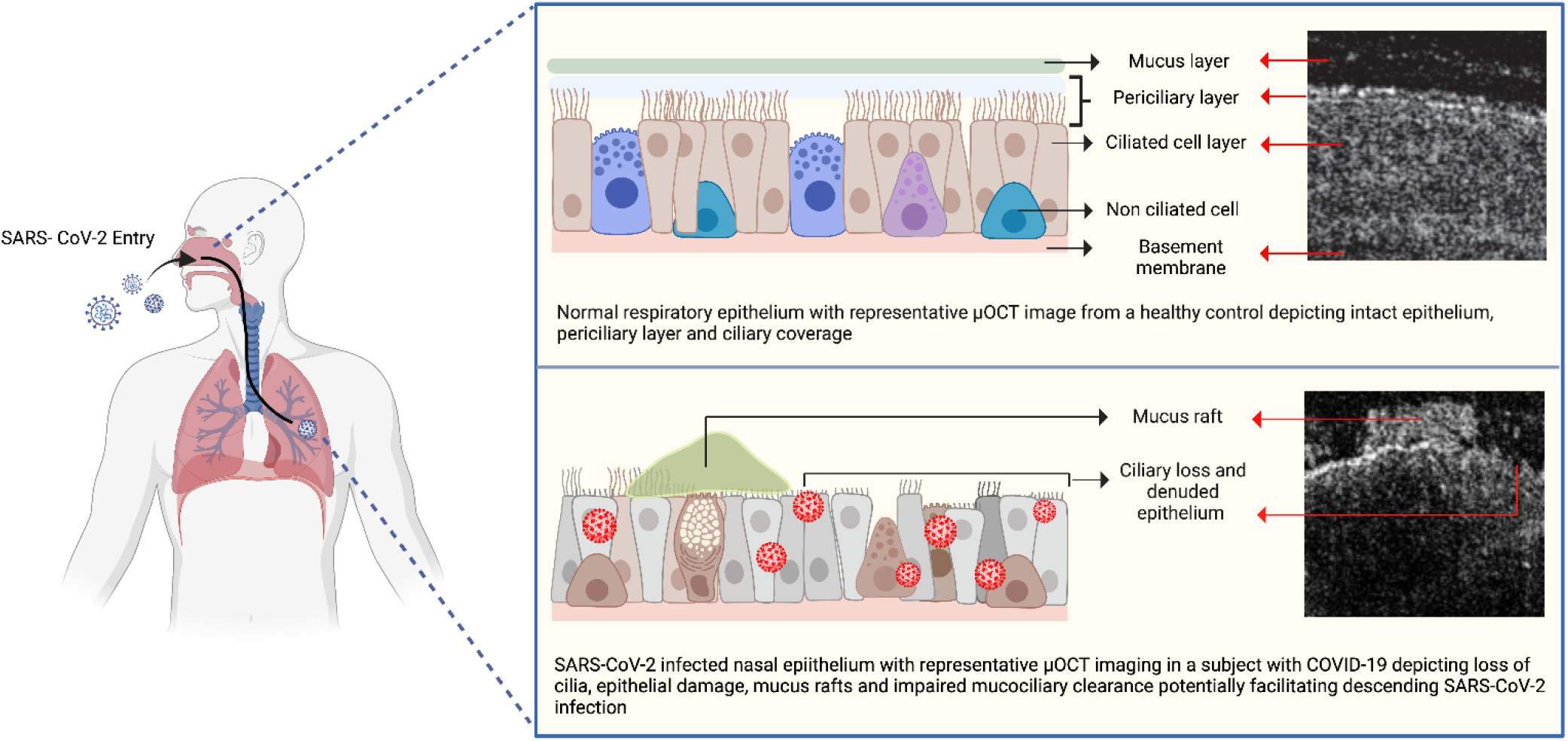

## INTRODUCTION

Selective tropism for respiratory transmission of severe acute respiratory syndrome coronavirus 2 (SARS-CoV-2), the causative agent of the global pandemic of coronavirus disease 2019 (COVID-19), is attributed to high angiotensin-converting enzyme 2 (ACE2) expression, the functional receptor for the virus, in ciliated respiratory epithelial cells. (1, 2) The entry of the virus is likely dependent on the initial viral load, and the rapid spread of COVID-19 may be attributable to the efficient transmissibility of SARS-CoV-2 involving the ciliated cells of the upper airway. (3) Clinical presentation of COVID-19 illness from asymptomatic to more severe pneumonia and respiratory failure occurs through a variety of mechanisms including cytopathic cellular effects and viral mediated cytokine responses. (4) Molecular evidence in several single cell RNA sequencing studies have implicated the upper respiratory tract to be the gateway for replication and transmission for the SARS-CoV-2 virus (1, 2, 5, 6) thus necessitating a better discernment of the functional pathophysiology of the ciliated nasal epithelium.

Several *in vitro* and *in vivo* studies have demonstrated proclivity for SARS-CoV-2 virus to ciliated cells, evidenced by prominent ciliary abnormalities (7–9) and impaired mucociliary clearance (10, 11) in these model systems. Perhaps consequently, hyper viscous mucus in bronchoalveolar lavage (BAL) fluid (12–14) and in the airways (15, 16) of COVID-19 patients have been reported, suggesting the clinical importance of these events. Despite this, our understanding of the early pathogenesis of COVID-19 on the nasal epithelium in human subjects remains incomplete, particularly in regards to early physiologic events and their functional consequences on ciliary beating, mucus properties, mucosal inflammation, and the mucociliary transport apparatus.

The study of airway pathophysiology in humans has been limited by the absence of an *in vivo* imaging technology that is capable of visualizing the respiratory mucosal apparatus at the subcellular level. Addressing this gap, we developed intranasal micro optical coherence tomography (µOCT) and applied it in patients with cystic fibrosis (CF) to characterize the functional microanatomy, identifying diminished airway surface liquid (ASL) and periciliary layer depths (PCL), and delayed mucociliary transport rate (MCT), among others key metrics of epithelial function, helping ascertain the underlying pathophysiology of the disorder. (17) Here, we devised a method to safely isolate individuals with symptomatic COVID-19 and used intranasal µOCT imaging to characterize the functional consequences of SARS-CoV-2 infection on mucociliary homeostasis. Our results demonstrate profound ciliary loss in the nasal epithelium even in outpatient subjects with mild disease and sheds light on mechanisms that could foster disease progression and predispose patients to secondary infections that commonly complicates and prolongs COVID-19 related illness.

## RESULTS

### Subjects

36 subjects who were known COVID-19 positive either by rapid antigen screen or RT-PCR testing, and above the age of 18 were screened between October 2020-April 2021. Sixteen subjects were enrolled for µOCT imaging. Thirteen subjects were positive by RT-PCR at the time of imaging and included in the analysis. Three subjects tested RT-PCR negative, with no detectable viral loads for SARS-CoV-2, had a history of partial vaccination (single dose of BNT162b2 vaccine (Pfizer-BioNTech)), and were hence excluded from the final blinded µOCT analysis. For the control group, owing to the difficulty in recruiting normal subjects during the pandemic, seven subjects who were imaged between May 2018-October 2020 were included in the blinded µOCT analysis in addition to the 1 subject enrolled during the COVID-19 cohort enrollment period. (Figure 1, consort diagram).

**Figure 1.**
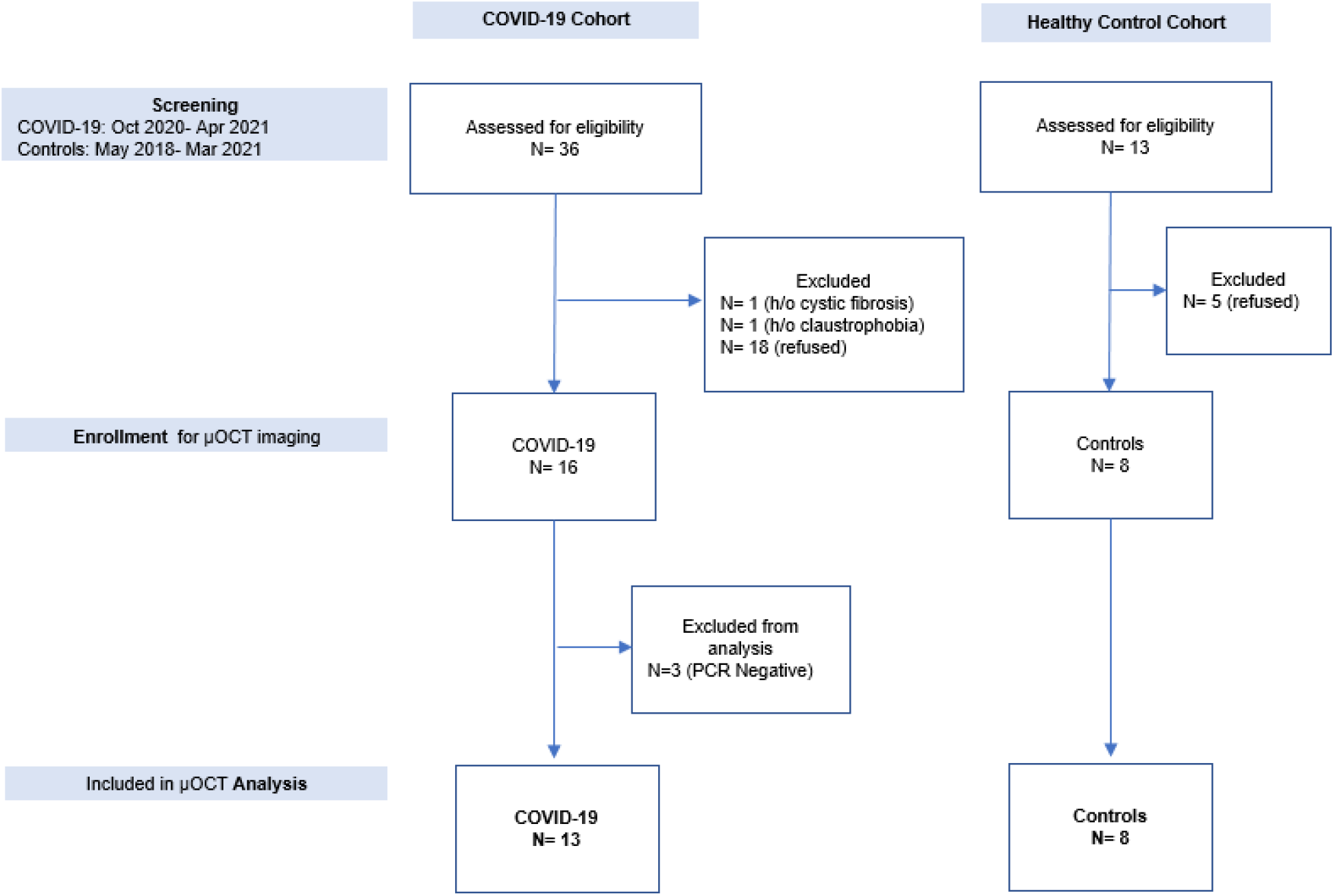
Consort diagram.

There was no significant difference in age or sex distribution between the COVID-19 and control groups at baseline (Table 1). The mean age of subjects in the COVID-19 group was 35 years, 54% were female; 69% were White, 23% Black and 8% Asian. All of the subjects presented with symptoms of mild COVID-19, and scored 1-2 (mild, ambulatory) on the WHO Ordinal scale for COVID-19 disease severity. Obesity was the only known risk factor in the COVID-19 group. Fatigue was reported in all COVID-19 subjects, followed by fever and chills, dyspnea and cough in 92% each. Three subjects (23%) of the subjects had nausea, vomiting and diarrhea. The median viral load was 4.28 log_10_ copies per ml. Three subjects (23%) had a viral load of ≥10^4^ copies per ml (Table 2). Follow-up history of subjects with COVID-19 at the end of 21 days from the day of µOCT imaging did not reveal worsening symptom burden or progression of disease course including need for supplemental oxygen, or hospitalization.

**Table 1.** Demographic and clinical baseline characteristics.

| **Subject Characteristics** | **COVID-19 (N= 13)** | **Healthy Controls (N=8)** |
| --- | --- | --- |
| Age (years ± SD) | 35 ± 8.4 | 31 ± 6.9 |
| Sex (female, N (%)) | 7 (54%) | 4 (50%) |
| Race: White/ Black/ Asian (N) | 9/3/1 | 6/2/0 |
| Smoking status (N) | 0 | 0 |
| BMI (kg/m<sup>2</sup>) * | 41.2 ± 18.1 | - |
| Healthcare workers (N) | 4 | 0 |
| Illness duration (days ± SD) ^ | 10.15 ± 2.2 | N/A |

\*BMI, the body-mass index is weight in kilograms divided by the square of the height in meters.
^Illness duration was defined as days from symptom onset to imaging.

**Table 2.** Presenting symptoms and severity indicators of COVID-19 cohort.

|  |  |
| --- | --- |
| **Symptoms (N, (%))** |  |
| Fatigue | 13 (100%) |
| Fever and chills | 12 (92%) |
| Cough | 12 (92%) |
| Dyspnea | 12 (92%) |
| Loss of smell and taste | 10 (77%) |
| Headache | 9 (69%) |
| Nasal congestion | 9 (69%) |
| Sore throat | 5 (38%) |
| Myalgias | 5 (38%) |
| Nausea and vomiting | 3 (23%) |
| Diarrhea | 3 (23%) |
| **WHO Ordinal Scale for Disease Severity (range)** |  |
| 1 – Ambulatory, with no limitation of activities | 8 (62%) |
| 2 - Ambulatory, with some limitation of activities | 5 (38%) |
| **Co-existing conditions (N (%)) \*** |  |
| None | 10 (77%) |
| One | 3 (23%) |
| Two or more | 0 |
| **Viral load (mean, log<sub>10</sub> copies per mL)** | 4.28 |

\*Co-existing conditions included anxiety/ depression, migraine and hypertension in one subject each.

### Imaging procedure

µOCT imaging was conducted on all COVID-19 positive subjects within 14 days of symptom onset to coincide with the peak infectivity period (average 10.2 ± 2.2 days). The subjects underwent nasal imaging while seated in the COVID-19 isolation benchtop box with their head stabilized on an ophthalmologic head rest (Figure 2). The µOCT imaging procedure was well tolerated overall. 1 subject with COVID-19 experienced transient lightheadedness, sweating, and dizziness compatible with presyncope but had no alteration in vital signs, was self-limiting and did not affect completion of procedure. No adverse events were noted in the control group. No mucosal swelling or injury was observed during or immediately after the procedure in any of the test subjects.

**Figure 2.**
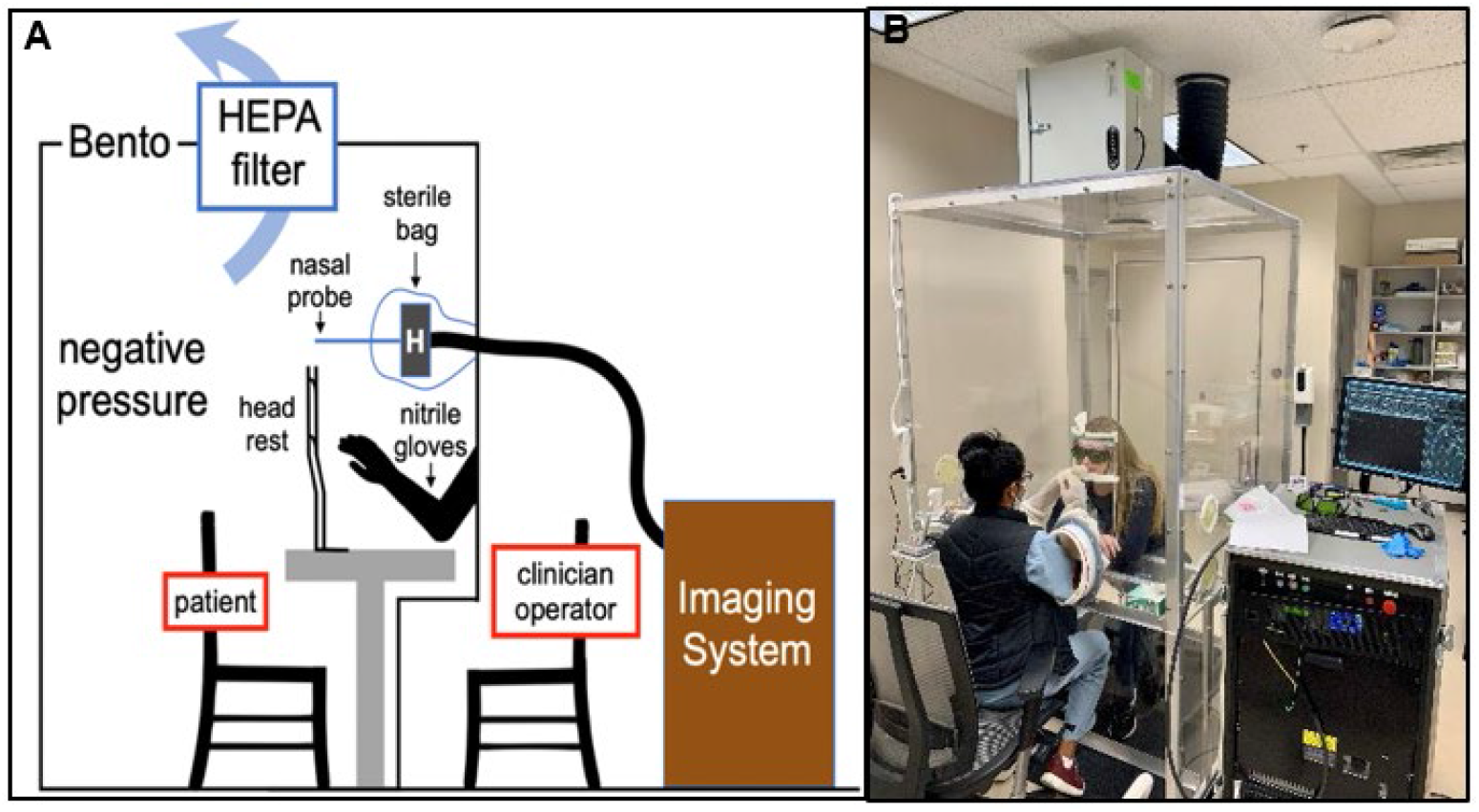
COVID-19 study subject isolation booth and demonstration photograph. The personal protection booth is a closed negative pressure HEPA filtration unit, with glove port and intercom communication access (A). Imaging was performed with the subject seated inside the box with chin rested on the ophthalmologic head rest, and operator interfacing with subject from the exterior through the glove ports (B).

### µOCT parameters

Micro-optical coherence tomography (μOCT) allows for visualization and quantitation of microanatomic parameters that characterize the function of the mucociliary clearance apparatus. Parameters that can be readily estimated include the periciliary liquid depth (PCL), cilia beat frequency (CBF), the area of active ciliary beating, termed percent ciliary coverage (pCC), and the mucociliary transport (MCT) rate. Quantitative metrics were extracted and analyzed by a team blinded to condition. Data was interpreted for COVID-19 subjects (n=13) in comparison to previously and concurrently imaged healthy controls (n=8). Of all 21 subjects imaged, the success rate of having at least 2 technical replicate values for each of the quantitative metrics were 100% for pCC, 62% CBF, 100% for PCL, and 43% for MCT. Success rates were lower for CBF (38% for COVID-19 vs. 100% for controls) and MCT (15% for COVID-19 vs. 87% for controls) metrics in COVID-19 infected individuals compared to healthy controls due to injury to the epithelium. In comparison to healthy controls and prior experience (17), µOCT imaging readily identified abnormalities in COVID-19 subjects including denuded epithelium with loss of ciliated cell function, presence of mucus rafts, and increased inflammatory cell counts suggesting that individuals with mild but symptomatic COVID-19 exhibit functional abnormalities of the respiratory mucosa that can be captured by µOCT (Figure 3, Figure S2).

**Figure 3.**
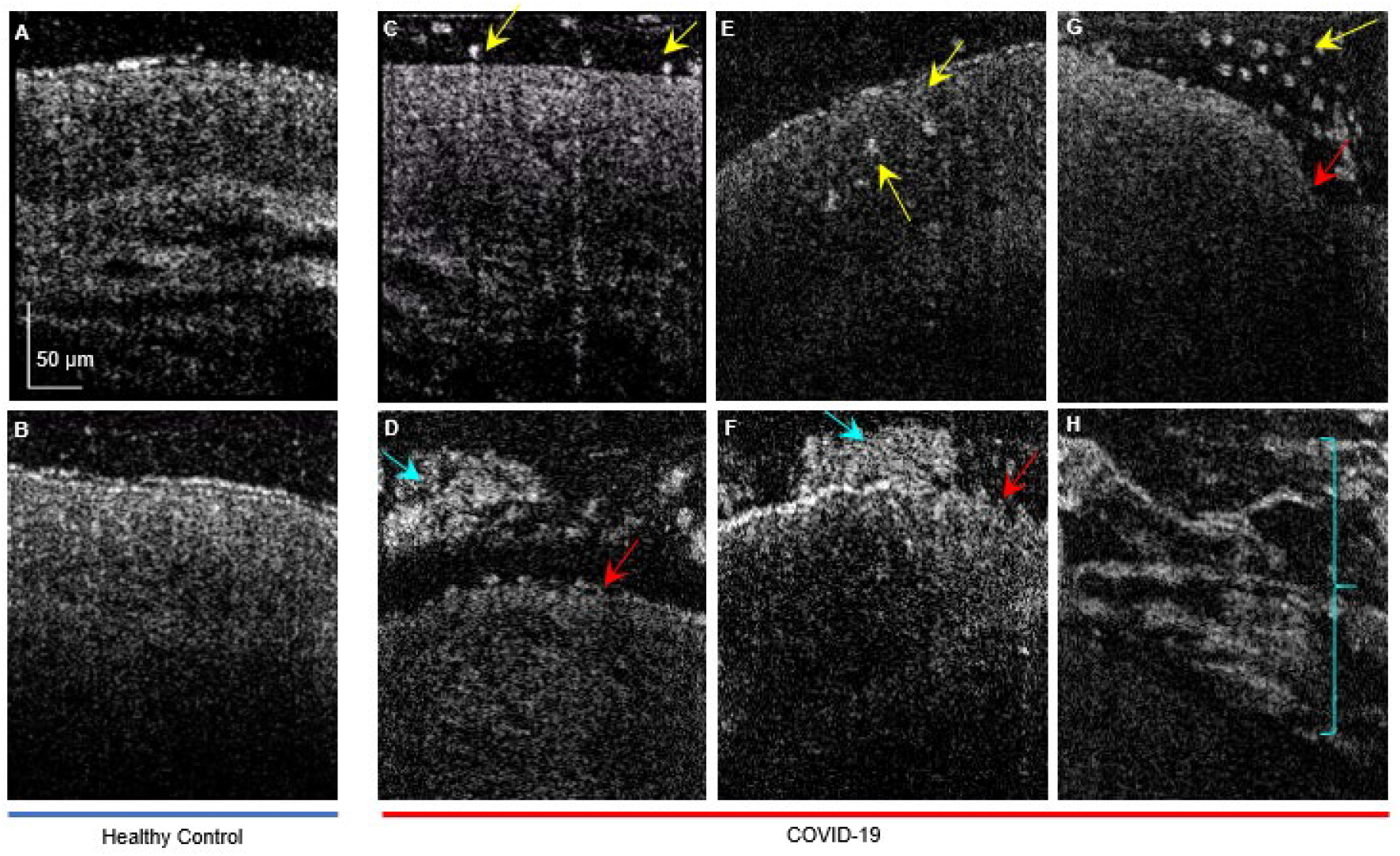
µOCT imaging demonstrates multiple abnormalities in subjects with COVID-19. Healthy controls (A-B) with uniform PCL layer, preserved epithelium and absence of mucus tufts or inflammatory cells versus COVID-19 subjects (C-H) with readily apparent mucus rafts (blue arrows), denuded epithelium and loss of ciliary coverage (red arrows), and increased inflammatory cells (yellow arrows).

#### COVID-19 leads to severe loss of ciliary coverage in the nasal epithelium and reduces ciliary beat frequency

Unlike patients with cystic fibrosis, no meaningful difference was found for PCL depth between the healthy controls and COVID-19 subjects indicating the absence of a discernable defect in airway hydration in the COVID-19 cohort (Figure 4). In contrast, subjects with COVID-19 had severely reduced ciliary coverage (pCC, 57.4 ± 36.11% healthy controls vs 18.1 ± 23.62% COVID-19, *P*= 0.009). Although high viral loads have been associated with increased mortality, growing evidence suggests that initial viral loads may not always correlate with disease severity.(18, 19) Consistent with this, 23% of the COVID-19 cohort had significantly high viral titers at the time of imaging, yet there was no direct correlation between pCC and viral load at the time of imaging (R^2^ =0.17). Similarly, pCC was not affected by the duration of illness (R^2^ <0.01). (Figure S1)

**Figure 4.**
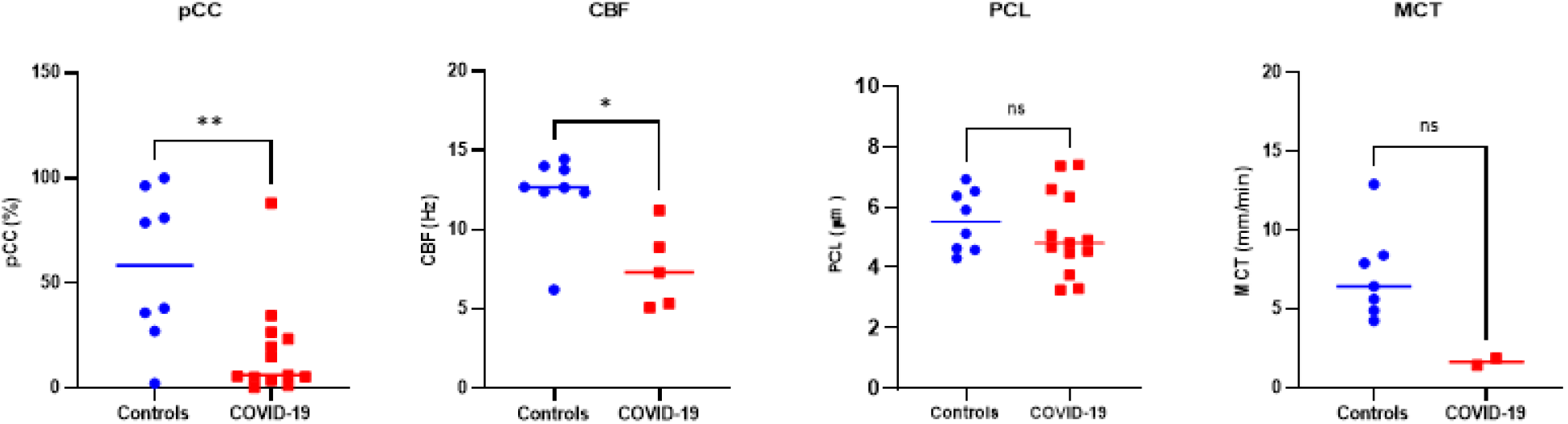
Functional microanatomical measurements are abnormal in COVID-19. Scatter plots of percent ciliary coverage (pCC), ciliary beat frequency (CBF), periciliary layer depth (PCL) and mucociliary transport rate (MCT) measurements of healthy controls (blue) in comparison to COVID-19 (red) subjects. Each data point represents the mean measurement for each individual. Bars are means and 95% CI. Comparison of data by Mann-Whitney test, *P<0.05; **P<0.01.

SARS-CoV-2 infection in experimental animals and *in vitro* is known to cause shortening and misshapen ciliary structure, thus contributing to alterations in function (10, 11), but this has not yet been assessed in symptomatic COVID-19 patients. µOCT demonstrated ciliary beat frequency was severely diminished in COVID-19 patients compared to controls (12.32 ± 2.58 Hz healthy controls vs 7.57 ± 2.56 Hz COVID-19, *P*= 0.011). To further evaluate ciliary function and examine residual ciliary activity, we performed qualitative waveform analysis as previously reported in excised rat trachea and cell culture (20) for the first time in human subjects (Figure 5). In healthy subjects, representative waveforms of cilia detected by our automated algorithm showed regular, rhythmic patterns of relatively consistent frequency and amplitude. In comparison to controls, representative ciliary beat waveforms in COVID-19 subjects showed striking differences, including irregular beat patterns with erratic amplitudes. This finding is consistent with data in hamsters where our group found that fewer motile cilia combined with abnormal ciliary motion of residual cilia contribute to delayed MCT in SARS-CoV-2 infection. (11)

**Figure 5.**
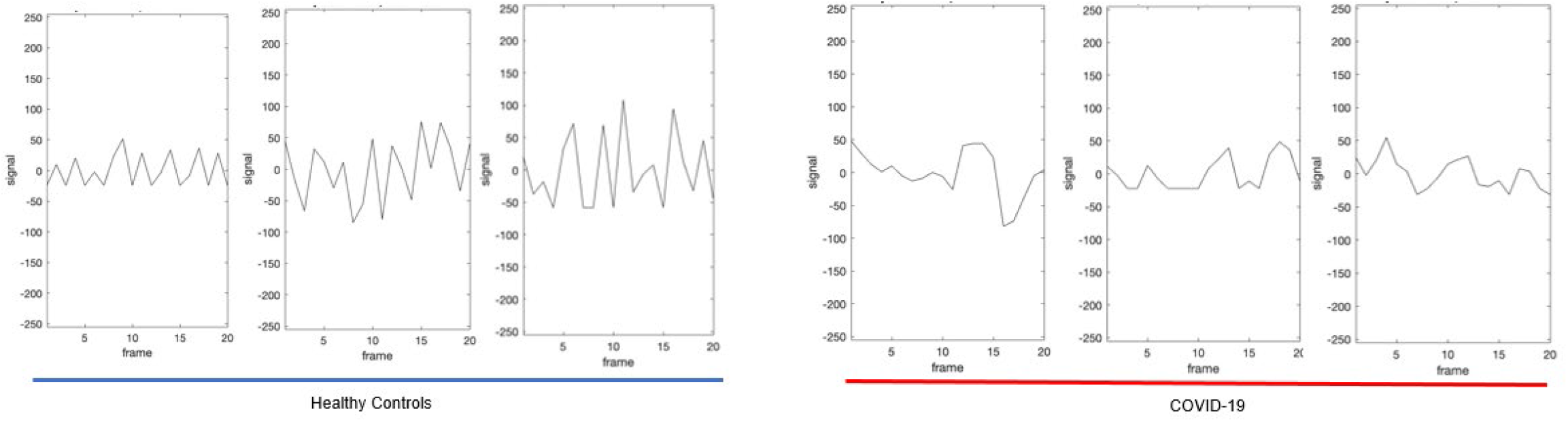
Representative ciliary beat frequency waveforms are abnormal in COVID-19. In comparison to healthy controls (left), who exhibited relatively consistent amplitude and frequency, representative waveform analysis of detected cilia in COVID-19 infected subjects (right) show irregular beat patterns with erratic amplitude.

Due to the severity of the abnormalities in the mucociliary transport apparatus, and frequent areas where the epithelium had complete absence of cilia, it was challenging to ascribe the motion of motile particles to mucociliary transport in majority of the COVID-19 subjects. In subjects where quantification was possible, mucociliary transport rates showed a pronounced reduction in the COVID-19 cohort as compared to controls, although this was not statistically significant (7.2 ± 2.9 mm/min healthy controls, n=7 vs

## DISCUSSION

Structural integrity and coordinated movement of the cilia are imperative for optimal mucociliary clearance, the primary innate defense mechanism of the respiratory tract.(21) Ciliary dysfunction has been associated with various disorders, including as the primary abnormality in primary ciliary dyskinesia and secondary abnormalities in cystic fibrosis, chronic obstructive pulmonary disease, and chronic rhinosinusitis.(22) In SARS-CoV-2 infection, infection of ciliated respiratory cells is prominent, but the functional consequences on the human nasal epithelium have not yet been defined. In the first rigorous reporting of *in vivo* real time evidence of ciliary dysfunction in patients with acute COVID-19, we found prominent and unique changes to the functional microanatomy of the nasal epithelium. Most importantly, we observed extensive loss of motile cilia, and this was accompanied by derangements in residual ciliary beating and loss of coordinated motion. Prominent cellular inflammation of the overlying mucus, with adherent mucus rafts were also apparent. These features are likely to contribute to delayed mucociliary clearance that could have important consequences on downstream pathogenesis, including removal of viral particles and protection against secondary infections of the respiratory tract. Of note, findings in COVID-19 contrasted with µOCT imaging of patients with CF where significantly reduced MCT rates were observed, but in relation to depletion of periciliary liquid layer and the presence of hyper viscous mucus. (17) Our results could be relevant to other respiratory viruses, as virus-mediated deciliation has been reported previously with coronavirus infections (23), and inflammatory cytokine mediated decreased ciliary activity and direct toxin mediated ciliary damage have been postulated in infections such as influenza and pertussis. (24–26)

Prolonged nasal mucociliary clearance times have been demonstrated in patients with mild COVID-19 using nasal saccharin testing, however these findings have been inconsistent depending on the duration of symptoms and the physiological cause has not been defined. (27, 28) Here we provide the first real time evidence for SARS-CoV-2’s tropism for ciliated cells in the nasal epithelium in patients, and its consequences on mucociliary transport function. These results are consistent with several *in vitro* and *in vivo* studies that have described ciliary injury as a common pathological feature in COVID-19 models. These characteristics have included prominent infection of the ciliated cells, in addition to disruption of cilia synchronicity and cilium shrinkage with decreased ciliary beating demonstrated in differentiated human tracheobronchial epithelial (HTBE) cells (7, 10, 29), nasal ciliated cells from human and non-human primates (5), Syrian hamsters (11, 30, 31), and SARS-CoV-2 infected human nasal organoids. (32) In addition, diminished mucociliary clearance by SARS-CoV-2 has been demonstrated in HTBE cells by tracking polystyrene beads (10) and in Syrian hamsters by µOCT imaging. (11)

By intention, due to requirements for conducting safe and effective nasal imaging under isolation conditions, our disease cohort had mild COVID-19 illness. Thus, we principally enrolled a population who were young and without substantial comorbidities at baseline. Obesity was the only known risk factor for severe COVID-19 illness (33) in this cohort. Despite this, we found severe ciliated cell abnormalities and functional deficiencies in the mucociliary apparatus even in early, mild COVID-19 illness when imaged within the peak infectivity period. More specifically, we observed a 40% decrement in pCC highlighting profound loss of ciliary coverage, a finding that was more extensive than that found in CF even though epithelial remodeling due to chronic infection is prominent in that disorder.(17) Residual cilia function was also abnormal, with a 48% decrease in CBF and abnormal waveforms of remaining ciliary tufts. Ciliated cell loss in our study was found to be independent of the viral loads and duration of illness and is reflective perhaps, of additional factors including innate immunity, genetic and altered host inflammatory responses as being responsible for disease progression. It is noteworthy that viral loads have failed to have a consistent prognostic potential for the severity of illness in several cohorts, (34–36) hence highlighting the possibility that functional parameters may hold more promise for disease stratification.

While we attempted to more comprehensively examine MCT, a key marker of host defense, the inability to discern motile particles in the background of severe ciliary loss reduced our ability to accurately measure MCT in many cases. In µOCT studies of excised trachea in Syrian hamsters, we documented severe decrements in MCT that accompanied ciliary loss; these studies benefited from laboratory conditions where MCT could be more readily measured. (11) Mucus stasis, as observed in our study by elucidation of adherent mucus rafts and other features of pathologic mucus, is known to increase the risk of secondary infections. The consequent breakdown of primary defense mechanism and infectivity gradient from the upper to lower respiratory tract may be responsible for the cascade of progression from mild disease to more severe pneumonia and acute respiratory distress syndrome (ARDS), (37) while portending a risk for bacterial or fungal superinfections occurring in up to one third of COVID-19 patients. (38–43). Notably, excessive sputum production has been described in up to 20% of patients with COVID-19, (44) and over one-fourth of critical ill COVID-19 patients are known to have secondary infections. (45, 46) Additionally, autopsy studies in patients with severe COVID-19 have shown extensive mononuclear epithelial infiltration and mucus plugging in the affected lobes of the lung, (47, 48) further emphasizing the role of adequate ciliary health and optimal mucociliary clearance.

Our findings are limited by small sample size, and while we captured a variety of functional deficits in relatively healthy subjects with mild disease, their course was self-limited, thus restricting our ability to discern the implications of the functional abnormalities on the risks of future complications. Since serial imaging through the disease course was not feasible at the time of this initial study, we are unable to comment of the longitudinal progression of functional and microanatomical changes. The study was conducted early in the pandemic, before multiple variant surges and prior to the advent of widespread vaccination. Future work could examine the effects of more common variants such as omicron where nasal replication is even more extensive than the founding variants. MCT measurements require visualization of ciliated epithelium to define trackable mucus particles, limiting the number of cases where this could be accurately measured. In addition, imaging through the isolation box interface may have contributed to decreased image stability, thus reducing optimal data extraction.

In summary, our study highlights the use of μOCT imaging in investigation of early pathogenic mechanisms of COVID-19. Notably, despite the ambulatory status of the subjects evaluated here, ciliary abnormalities were prominent. As asymptomatic individuals are also known to propagate the virus, it seems highly likely that substantial ciliated cell injury may also be occurring in these individuals, hence underscoring the importance of targeting ciliated cells for early mitigation of disease transmission. Therapies directed towards improving function of the mucociliary apparatus deserve further investigation as a treatment modality to improve respiratory function, abrogate descending infection, and prevent secondary infections.

## MATERIALS AND METHODS

### Study design

#### Study subjects

Symptomatic outpatients with the ability to provide informed consent and suspected COVID-19 who presented to the University of Alabama at Birmingham (UAB) COVID clinics were recruited within 7 days of symptom onset. Inclusion criteria included age 18 years and older, and confirmed COVID-19 as determined by either RT-PCR or rapid antigen for SARS-CoV-2. Subjects were excluded if they were unwilling to comply with institutional policies for personal protective equipment use, had moderate to severe COVID-19 illness requiring hospitalization, need for supplemental oxygen therapy or continuous positive airway pressure for respiratory insufficiency, were pregnant, or had history of preexisting lung disease or major sinus surgery that altered nasal anatomy and precluded imaging of the nares.

#### Clinical data collection

Detailed clinical history including demographic factors, COVID-19 symptoms, COVID-19 exposure and vaccination history, past medical history, smoking history, and medication use. A COVID-19 risk factor questionnaire was administered. Abbreviated physical exam and body weight were conducted. Long-term follow up, including clinical outcomes at 21 days, and any cause for hospitalization was collected.

### Procedures

#### Design of the full-body personal protection booth

Since intranasal µOCT is potentially an aerosol-generating procedure, a custom, negative-pressure full-body personal protection booth (Figure 2) was designed and fabricated to isolate the study subjects from the study staff. The booth was made compatible with the intranasal µOCT imaging system and its operation. The body of the booth was 4 x 4 x 7 ft and the booth contained a HEPA filter and air suction unit that isolated the study subject inside the booth. All walls of the booth were transparent to allow visualization of the live µOCT video during an imaging session and to facilitate communication. An intercom was also installed to transmit audio between the person inside and outside of the booth. A horizontal platform was integrated onto the front wall of the booth to aid the clinician to achieve a good and stable placement of the intranasal probe. A headrest, important stabilizing the study subject during imaging, was installed on the horizontal platform. Leg space was integrated onto the front wall of the booth to allow both the clinical and the imaged subject to sit across each other during the imaging procedure. A flapped wall port was used to enable the insertion of the fiber-optic µOCT probe into the booth. The µOCT imaging console was external to the subject isolation booth.

#### Infection prevention and imaging procedure

To protect study subjects and staff, research subjects underwent nasal imaging while seated in the custom designed personal protection booth (Figure. 2). This system was used to isolate the study subject from the study staff and the µOCT equipment via negative pressure HEPA filtration system certified by MGH COVID and UAB Infection Control. The study staff clinician imaging the patient performed the µOCT imaging procedure from outside the booth by manipulating the hand-held piece and nasal µOCT probe from the outside in, using sealed glove ports integrated in the booth. The hand-held piece inside the booth was covered with a sterile plastic sheath which was disposed following each procedure. Following inspection of the nares with an illuminating rhinoscope, the µOCT probe was maneuvered into the inferior meatus region while visualizing real time images and acquiring data from approximately five discrete sites at the turbinate and floor of each nare. The interior of the booth was disinfected after each study subject’s imaging session, followed by a one-hour decontamination period with continued highly efficient particulate air (HEPA) filtered flow. All study staff were equipped with respiratory enhanced personal protective equipment for medical providers as an additional precaution.

#### Viral load quantification by RT-PCR

Immediately following the imaging procedure and nasal inspection, mid-turbinate nasal swabs were collected for RT-PCR to confirm SARS-CoV-2 infection. The samples were obtained using disposable sampling swabs manufactured by Luxus Lebenswelt GMBH (Beijing NaGene Diagnosis Reagent Co., Ltd) and transported using Thermo Scientific™MicroTest™ M4RT 3mL tubes, containing gelatin, gentamicin, and amphotericin B. Nasopharyngeal swab samples were evaluated by RT-PCR in the CAP-accredited UAB Fungal Reference Laboratory as indicated below.

The Maxwell^®^ RSC Viral Total Nucleic Acid Purification Kit (cat. # ASB1330, Promega Corporation, Maddison, WI) was used to isolate RNA as instructed by the manufacturer. Briefly, 300µl of the sample was mixed by vortexing for 10 seconds with 330 µl of freshly made mixture of lysis buffer and proteinase K solution at 10:1 ratio. Samples were incubated at room temperature for 10min then at 56°C for another 10min, and extracted utilizing the Maxwell^®^ RSC 48 Instrument (Promega Corporation). qRT^2^-PCR was performed utilizing the QuantStudio™ 5 Real-Time PCR System (Fisher) and the following primers/probes: (forward 5’-GAC CCC AAA ATC AGC GAA AT-3’ and reverse 5’-TCT GGT TAC TAC TGC CAG TTG AAT CTG-3’) and probe (/SFAM/ACC CCG CAT TAC GTT TGG TGG ACC/BHQ_1) targeting the nucleocapsid (N) gene. AccuPlex™ SARS-CoV-2 Reference Material (cat. # 0505-0126, SeraCare Life Sciences, Inc., Milford, MA) were processed in parallel with samples and used as standards for genomic viral load quantitation.

#### μOCT technology

The intranasal μOCT technology has been described in detail in previous publications. (17) In brief, the device is based on the principles of spectrometer-based spectral-domain OCT. Broadband light from a super continuum laser (SuperK Extreme OCT, NKT Photonics) is first filtered to achieve a Gaussian-like spectral shape. A beam splitter then directs a portion of the light to a custom-built fiber-optic intranasal probe, which accomplishes common-path interferometry at the distal end. With the side-viewing intranasal μOCT probe, an annular imaging beam was scanned over a lateral distance of 350 μm at a rate of 40 Hz. This rapid scanning action occurred within a sterile, single-use sheath that was fitted over the fiber-optic core and fixed onto the handle unit. The 2.4 mm diameter, semi-flexible sheath has a transparent imaging window that enables the beam to be transmitted and focused onto the tissue. This μOCT technology achieves a transverse and axial resolution of 4 µm and 1.3 µm in tissue (estimated refractive index, n = 1.4) respectively. The high-resolution cross-sectional images recorded at video rate with this technology were critical for enabling the measurement of discrete hydration layers of the epithelium, as well as for the characterization of sub-cellular features. Furthermore, the performance of this technology allowed dynamic events such as mucociliary transport and cilia beating to be captured. Apart from the intranasal probe and the attached handle unit, the entire optical system was installed on a compact wheeled cart, which facilitated its use in a clinical setting.

#### Image acquisition, stabilization, and analysis

All data was de-identified and stored for interpretation by a team blinded to the study visits. Data was compared to previously imaged historical age- and sex-matched control group. A total of 20 distinct anatomical locations from both nares were imaged. On each side of the naris, µOCT data were acquired from both the turbinate and floor of the nasal meatus. This imaging procedure resulted in an average of 29.6 30-second µOCT videos recorded from each subject. Each of the videos were first corrected for lateral scan non-uniformity before being scaled to achieve isotropic pixel-to-length representation. As all data were acquired on unsedated subjects by manual placement of the probe in the subject, stabilization of the µOCT videos had to be performed before they could be further analyzed. This was accomplished by first computing the local correlation maxima between two adjacent frames within a running 256 µm square window. The lateral shifts corresponding to the global correlation maxima were then used to register the frames. The level above which the correlation maxima exceeded the standard deviation was used to benchmark how well the in-plane registration worked which was then used to automatically select sections of the µOCT videos that were stable over a minimum of 20 frames or longer for further analysis of CBF and MCT. The method with which CBF, MCT, and pCC were extracted are described in previous publications. (17) In brief, CBF were computed by Fourier analysis of the µOCT signals modulated by beating action of the cilia. To characterize ciliary waveforms of detected cilia, the m-mode µOCT amplitude of the ciliary signal generated by ciliary motion were plotted for selected sub-regions of interest. MCT measurements were determined by tracking native mucus particles within 70 µm of the epithelial surface. To ensure that readings were taken from particles that were transported by native ciliary motion, those that mirrored the trajectory of the probe were deemed to be artificially influenced by the motion of probe and were excluded in the analysis. pCC was determined by calculating the percentage of the epithelial surface that was covered in motile cilia in a randomly selected frame within each acquired video.

#### Statistical analysis

Means were computed for each subject when at least two measures for each of the metrics were valid; mean values were used for subsequent cohort analysis. Data statistics are presented as means (SD), unless indicated otherwise. Differences in each metric between COVID-19 and normal cohorts were compared cross-sectionally with either unpaired *t* test with Welch’s correction or Mann-Whitney test, depending on whether the distribution of the measurements passed the Shapiro-Wilk normality test. Two-tailed *P* values are reported. P values of < 0.05 were considered statistically significant. Linear regression and the Pearson correlation coefficient (r) was for correlation analysis. Statistical analyses were performed using GraphPad Prism (version 9.1.0, GraphPad Software, San Diego, California USA).

#### Study approval

The study was approved by The Partners Institutional Review Board (IRB), Massachusetts General Hospital (protocol #2016P000272, isolation booth approved in AME #21, Aug 2020) and the University of Alabama at Birmingham IRB (protocol #F160125001). All subjects provided written informed consents prior to participation.

## Author contributions

KV, HML, GMS, GJT and SMR developed the study design

KV, GMS, HYH, JDW, KM, KBS and SMR participated in experimental conduct and acquisition of data

KV, HML, AB, CMFP, GMS, GJT and SMR performed data analysis

SML, DBM and HMP performed virologic testing and analysis

KRO, PC, SF, DM, GJT and SMR participated in the design and development of the personal protection booth.

KV, HML, QL and SMR prepared the manuscript.

All authors provided necessary revisions and approved a final draft prior to submission.

## Acknowledgements

This study was funded by the NIH (P30DK072482 (SMR), R35 HL135816 (SMR) and R35 HL135816-04S1(SMR), UL1TR003096 (UAB)), the Cystic Fibrosis Foundation (ROWE19RO, ROWE17XX1, TEARNE16XX0), institutional resources from the University of Alabama at Birmingham (COVID-19 Research Award) and funding from the Mike and Sue Hazard Family Foundation (GJT), and the Remondi Family Foundation (GJT).

Graphical abstract was created with biorender.com.

The authors are grateful to the participants enrolled in this study, and their families.

