## Supplemental Material for "COVID-19 Causes Ciliary Dysfunction as Demonstrated by Human Intranasal Micro-Optical Coherence Tomography Imaging"

**Figure S1. Correlation analysis between viral load, illness duration, and degree of functional ciliary expression.** (A) Correlation plots of percent ciliary coverage (pCC%) with viral loads (copies per mL) showed no direct correlation (r= 0.41). (B) Similarly, pCC% had no direct correlation with illness duration defined as days from symptom onset to time of imaging (r= 0.002).

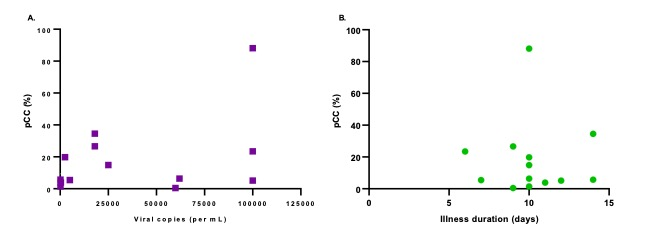

**Figure S2**. **µOCT imaging features apparent in COVID-19 illness.** Representative images from COVID-19 subjects with a variety of mucus accumulation patterns including multiple rafts, and thick adherent plaque-like accumulation (A-C), layers of mucus stranding (D-E), and mucus accumulation rich in inflammatory debris (F).

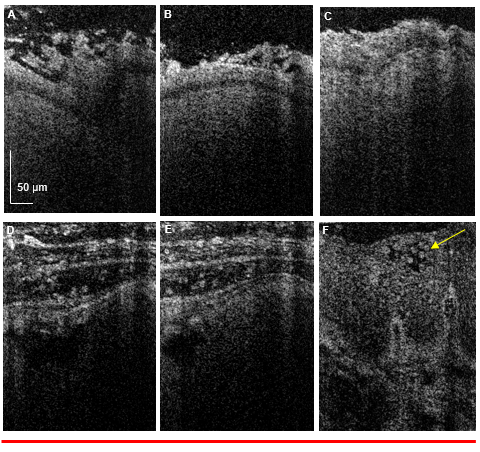
